# HDX-MS for Analysis of Protein Dynamics in Complex and Crowded Cellular Environments

**DOI:** 10.64898/2026.11.25.769177

**Authors:** Bindu Y Srinivasu, Shreya Ghosh, Tanvi S. Damerla, Thomas E. Wales

## Abstract

Hydrogen-deuterium exchange mass spectrometry (HDX-MS) offers potent insights into protein conformational dynamics and interactions but traditionally requires proteins to be purified to near homogeneity and to be removed from their native cellular context which is often challenging for unstable, untagged, or difficult-to-purify targets. Here we present “crude-HDX, crHDX” a workflow that we used to extend HDX-MS analysis progressively from purified proteins to unfractionated cell lysates and, ultimately, intact cells. Using KRAS as a model system, overexpressed protein was detected with sufficient sequence coverage and peptide redundancy for robust HDX-MS analyses directly within complex cellular backgrounds. By optimizing in-cell deuterium labeling, quenching, controlled cell disruption, proteolysis, and LC-MS conditions, we preserved deuterium signal while maintaining high peptide resolution. Comparison of KRAS across purified protein, lysate, and intact-cell environments revealed reproducible, time-dependent deuterium uptake with both conserved and environment-specific conformational features. To test whether crHDX can resolve protein-protein interactions in-cell, we co-expressed KRAS with the catalytic domain of SOS (SOS_cat_) and analyzed the complex directly in intact cells: SOS_cat_-induced perturbations in KRAS recapitulated key conformational HDX difference signatures seen with purified proteins while uncovering additional, cellular-context-dependent changes in HDX. Extension of the workflow to untagged KRAS and to the purification-challenging kinase GSK3β further demonstrates its generality beyond affinity-tagged, readily purifiable targets. Collectively, these results establish crHDX as a complementary strategy to conventional HDX-MS, enabling structural and interaction analyses of proteins with minimal purification and within increasingly physiologically relevant cellular environments.

## Introduction

Proteins are dynamic macromolecules that sample ensembles of conformational states rather than existing as immutable structures (1). Backbone dynamics are fundamental to protein function and regulate processes including molecular recognition, catalysis, allostery, signaling, protein-protein interactions, and ligand binding (2–4). Importantly, protein dynamics are strongly influenced by the surrounding molecular environment. Within the cell, any analyte protein is embedded in a densely populated environment that contains 100s of millions of macromolecules and other smaller molecules including inorganic ions in flux, many of which can influence protein conformational equilibria (5). Consequently, structural and dynamic properties determined for an isolated protein under simplified *in vitro* conditions may not fully represent the native conformational ensembles populated within a cellular environment (6).

Hydrogen-deuterium exchange (HDX) coupled with mass spectrometry (MS) is a widely accepted approach that is used to investigate protein higher-order structure and conformational dynamics (7–9). HDX-MS monitors the exchange of backbone amide hydrogens with deuterium following timed exposure of a protein or proteins to an excess of deuterium *via* a D₂O-containing buffer. The rate of deuterium incorporation in the peptide backbone is influenced by both amide hydrogen bonding strength and solvent accessibility revealing local structural dynamics; changes in HDX report on externally modulated alterations to protein conformation/dynamics (10, 11). Following deuterium exchange, the reaction is typically quenched to both low-pH and temperature, the protein then is proteolytically digested, and the resulting peptides are separated by HDX-optimized liquid chromatography and analyzed by mass spectrometry (12). Since its initial introduction, HDX-MS has therefore found broad applications in mapping protein-protein and protein-ligand interactions, investigating allosteric regulation, characterizing protein folding and conformational transitions, and studying protein systems that may not be ammenable to other structural approaches (13–15). A major strength of HDX-MS is its ability to investigate structural dynamics of proteins both in solution and in the solid-state (16, 17) spanning a broad range of molecular sizes and can accommodate flexible, heterogeneous, and multidomain systems. Continued advances in sample handling (18, 19), chromatography (20, 21), mass spectrometry (22), and computational analysis (23–25) have substantially increased the sensitivity, reproducibility, and throughput of the technique. These developments have progressively expanded the utility of HDX-MS from relatively simple purified proteins toward increasingly challenging biological systems (26). Nevertheless, the vast majority of published conventional HDX-MS experiments continue to rely on proteins purified to near homogeneity prior to deuterium labeling (Figure 1). Explicit environmental control of protein purification offers several experimental advantages, such as protein concentration, ligand stoichiometry, nucleotide state, ionic composition, pH, and other experimental variables. This level of control facilitates a less complicated mechanistic interpretation of HDX measurements. However, the multiple chromatographic and sample-processing steps required during purification may disrupt weak or transient protein-protein interactions, remove endogenous ligands and metabolites, alter oligomeric equilibria, and potentially shift the conformational ensemble of the target protein from those populated within its native cellular environment (5).

**Figure 1.**
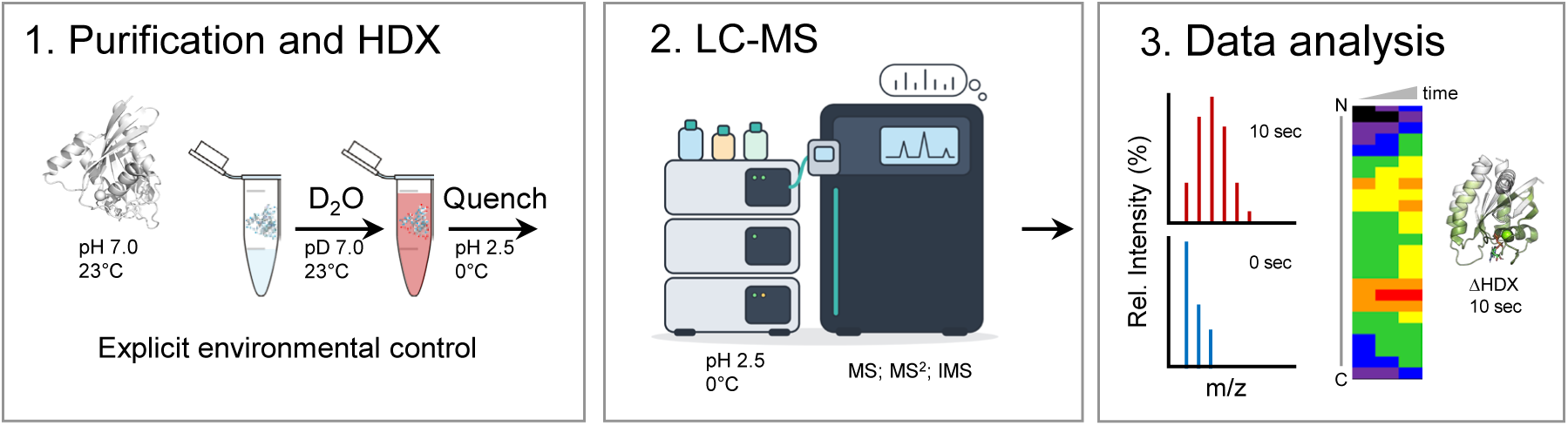
Conventional purification-dependent HDX-MS workflow. 1. Purified protein/protein:ligand/et. are subjected to D₂O labeling for defined exchange periods, followed by rapid quenching. 2. The resulting quench samples are digested under HDX-compatible conditions, and the resultant deuterated peptides are separated by liquid chromatography and analyzed by mass spectrometry (LC-MS). 3. The data are analyzed and visualized with various external programs.

With the above in mind, there has been a growing interest in extending structural mass spectrometry techniques towards increasingly complex biological samples while preserving physiologically relevant protein states and molecular interactions. Recent methodological advances have progressively extended HDX-MS from purified proteins to complex protein mixtures, in varying levels matrix complexity. These attempts depended on the use of affinity tags (27–29), protein thermal depletion (30), and either an overexpressed membrane protein of interest (31) or the study of hemoglobin in red blood cells by Sreekala *et.al.,* where hemoglobin comprises more than 90 % of cellular protein content (32). Extending such measurements to soluble intracellular proteins, particularly those for which no straightforward enrichment or fractionation strategy exists, remains considerably more challenging.

Moving from purified proteins to protein mixtures, unfractionated cell lysates, and ultimately intact cells therefore represents a progressive increase in both biological relevance and analytical complexity. Measurements performed in these environments have the potential to preserve features that are absent from purified-protein systems, including macromolecular crowding, endogenous binding partners, metabolites, ions, and transient molecular interactions. However, retaining cellular complexity rather than removing it through enrichment or fractionation places substantially greater demands on the analytical workflow. Target-derived peptides must be identified and quantitatively monitored against a large background of endogenous cellular peptides, while deuterium loss during post-labeling processing must be minimized. Additionally, complex mixtures introduce substantial analytical challenges for HDX-MS. In a conventional experiment with a purified protein, the number of peptides entering the LC-MS system is comparatively limited, allowing robust peptide identification and high sequence coverage. In a cell lysate or cellular sample, peptides derived from the target protein must be detected against a vastly larger background of endogenous peptides. Co-elution, ion suppression, overlapping isotope distributions, and competition for ionization can reduce the detectability of target peptides.

Here we present our work to develop a robust and adaptable methodological strategy that enables the HDX-MS analyses of soluble cytoplasmic proteins while preserving the cellular complexity during deuterium labeling; target-specific enrichment, fractionation, or other sample manipulations prior to traditional HDX LC-MS analysis are minimized. We have developed what we are calling a “crude HDX” (crHDX) workflow designed to minimize the requirements of extensive protein purification. This extends the reach of HDX-MS analysis from purified proteins to unfractionated cell lysates and ultimately within intact cellular environments. We first determined whether an overexpressed protein could be detected with sufficient sequence coverage and peptide redundancy within complex cellular protein backgrounds. We then optimized each step of the traditional HDX-MS experiment: deuterium labeling, quench, cell disruption, proteolytic digestion, and chromatographic separation to enable HDX measurements when labeling is performed directly within intact *E. coli* cells. The performance of our workflow was evaluated through comparison of HDX profiles across purified protein, unfractionated cell lysate, and intact-cell conditions of a well characterized protein of interest: KRAS 4b, which will be mentioned as KRAS. Together, this work establishes a methodological framework for performing HDX-MS with a progressively reduced dependence on conventional protein purification.

## Results and Discussion

### Development and optimization of crHDX in complex cellular environments: in-cell lysates

A vertical improvement in the reach of HDX-MS would be the ability to investigate protein conformational dynamics immediately following protein expression, while maintaining the protein in an environment that closely resembles its native cellular state. As such, we endeavored to establish an HDX-MS workflow capable of analyzing a protein of interest (POI) without requiring prior isolation to near homogeneity.

Traditionally, overexpressed proteins are purified as close to near homogeneity as possible prior to HDX-MS analysis, allowing their conformational dynamics to be examined in a simplified *in vitro* environment (Figure 1). Here, we sought both to establish and to optimize an HDX-MS workflow using unfractionated cell lysates, thereby minimizing the need for extensive purification of the over-expressed target protein. Under these conditions, the POI remains embedded within a complex mixture of host-cell proteins and other cellular components during deuterium labeling, better mimicking the features of the native intracellular environment, including macromolecular crowding and potential transient non-native protein-protein interactions. From a “back-of-the-envelope” calculation assuming that the average protein length in bacteria is 270 amino acids (33) that would mean there would be roughly 1.3 million peptides to separate (assuming 60-100 peptides per 100 amino acids of protein). All this from the nonspecific cleavages that are traditional in HDX-MS experiments. Addressing this level of analytical complexity requires substantial optimization of traditional HDX-MS workflows.

We sought to optimize 2 steps in the traditional HDX-MS experiment using a protein expression system with which we have substantial HDX-MS experience: KRAS (His₆-tagged 1-169) (34–37). First sample preparation and deuterium labeling, and second peptide level LC-MS separation and mass measurements were optimized for our POI, and the results are shown in Figure 2. In terms of sample preparation, we evaluated whether increasing on-column loads of clarified cell lysate yielded better sequence coverage of KRAS in the presence of increasing concentration of the host-cell proteome. Representative SDS-PAGE analysis of increasing amounts of clarified cell-lysate loaded onto the gel (2.5 - 15 μL) from a 4-hour post-induction expression of wild-type KRAS (residues 1-169) grown in *E.coli* BL21(DE3) cells is shown in Figure 2A. Purified KRAS is run alongside for reference. There is a prominent band between 20 and 25 KDa, that aligns with that of the band for purified KRAS and corresponds to the expected molecular weight of the overexpressed target protein (21.43 kDa). Additionally, there are bands for numerous endogenous cellular proteins readily detectable in each soluble fraction. Ultimately, we chose to use 5 μL of clarified cell lysate (based on theoretical calculations) as we were able to balance the robust overexpression of the target protein while also retaining the complexity of the surrounding host-cell proteome and total ion signal in the mass spectrometer. We next characterized the peptide composition of the digested lysate prior to deuterium labeling as in any traditional HDX-MS experiment through undeuterated control sample analyses. A representative chromatogram from an undeuterated control sample of lysate (5 µL) is shown in Figure 2B. Peptides in this chromatogram were generated in the following way: a 9-fold dilution of the lysate sample with buffer (50 mM HEPES, 150 mM NaCl, pH 7.8 in H_2_O) and equal volume of quench buffer (5 M guanidine hydrochloride, 0.8% formic acid, 2.5 % *n*-dodecyl-β-D-maltoside (DDM), 0.8 M Tris(2-carboxyethyl)phosphine (TCEP), and 1.5 mg *Aspergillus* protease, pH 2.3) was subjected to online digestion with an in-house prepared pepsin column (38, 39) without extended digestion *via* quench hold. The LC-MS analysis was performed using a chromatographic gradient of 5-35% acetonitrile in water (0.1 % formic acid) over 20-min and MS analysis using the Waters SELECT SERIES Cyclic IM QTOF mass spectrometer in HDMS mode. Analysis of the resulting LC-MS data using PLGS identified 94 peptides from KRAS totaling 80.47 % sequence coverage with an average peptide redundancy of 4.70 (Figure 2C). Both the broad sequence coverage and overlapping peptide redundancy demonstrated that an overexpressed target protein could be robustly detected and characterized without isolation to near homogeneity, even in the presence of a complex soluble host-cell protein background. In addition to KRAS, we identified 48 endogenous proteins representing diverse functional categories within the cell lysate (Figure 2D). Significantly, just under 70 % of these proteins were detected with greater than 60 % of peptide coverage. (Figure 2E). Not surprisingly, the majority of the proteins identified with significant coverage are all involved in protein production machinery. Together, these results demonstrate that our optimized LC-MS workflow to study a POI in complex matrices was able to identify not only the target protein directly within a crowded cell lysate environment using non-specific digestion, but also 48 additional host-cell proteins with high sequence coverage. Previous work in this area has not been as successful with host-cell proteins, and we believe that we can increase this number past 48 proteins with further optimization of the LC separation (31, 32).

**Figure 2.**
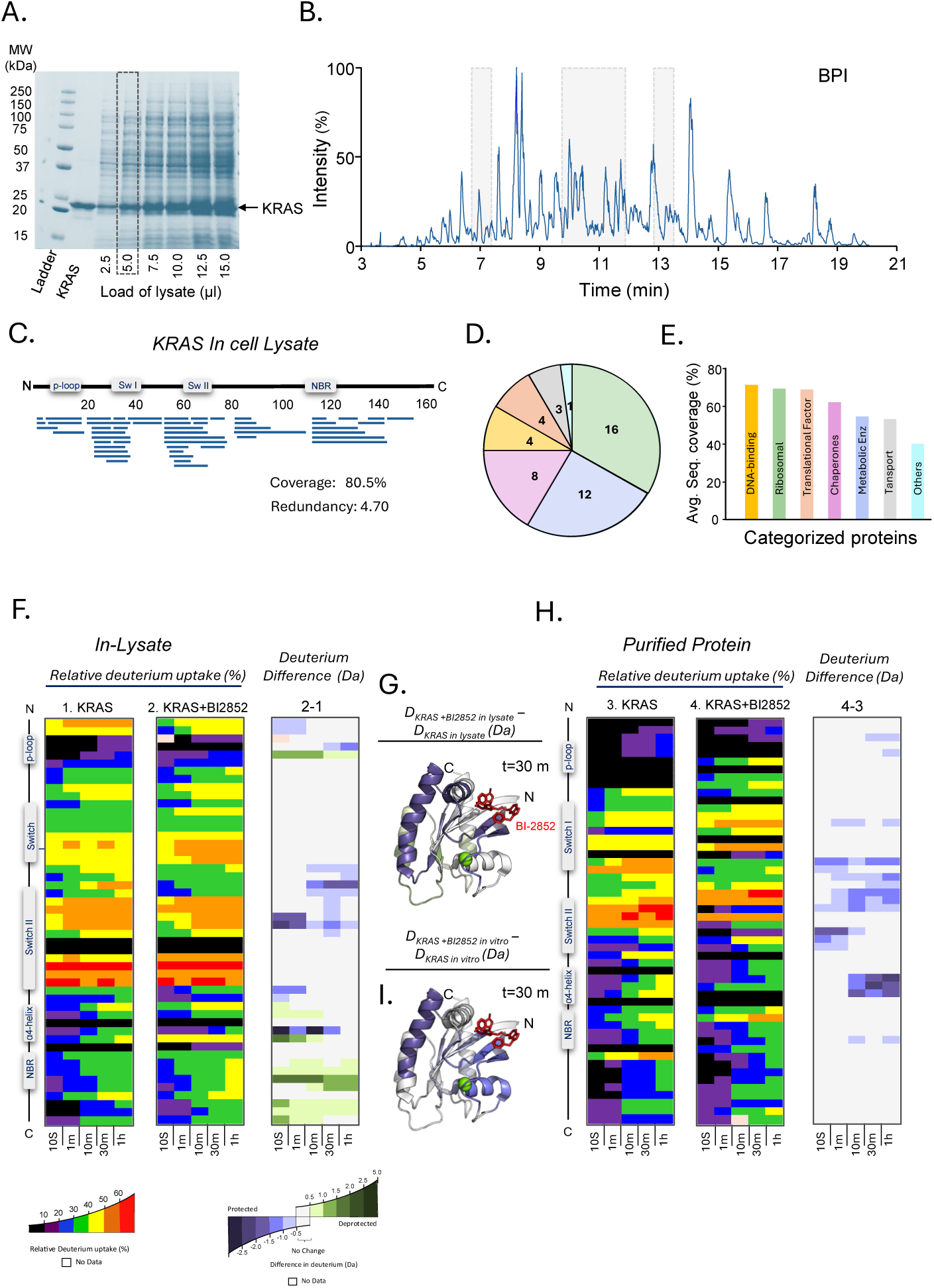
HDX-MS analysis of KRAS directly from a complex cell lysate: **(A)** SDS-PAGE analysis of the unfractionated *E. coli* cell lysate overexpressing KRAS. The highlighted region indicates the complex protein background used for subsequent analysis. **(B)** Representative LC-MS chromatographic profile of the proteolytically digested cell lysate; highlighted regions indicate time ranges where the bulk of KRAS peptides eluted. **(C)** Peptide coverage map for KRAS (excluding the His_6_ tag and linker) identified directly from the cell lysate. Blue bars represent individual peptides identified distributed across the KRAS sequence; major structural regions, including the p-loop, switch I, switch II, and nucleotide-binding region (NBR), are indicated. **(D-E)** Distribution of host cell proteins identified along with the POI, grouped according to their functional categories, demonstrating the diversity of proteins present in the sample. **(F–I)** HDX-MS analysis of KRAS directly within the complex cell-lysate background (F) compared to purified protein (H). Deuterium incorporation profiles are displayed as vertical peptide-level heat maps for a reduced set of peptides covering both the KRAS sequence and labeling time points (all HDX-MS data are included in the Supplemental excel file). Differences in HDX illustrates localized changes in KRAS conformational dynamics between the KRAS alone and KRAS in the presence of BI-2852. These changes in HDX are mapped onto the cartoon structure of KRAS PDB: 4OBE, (G) the effect on KRAS in the presence of BI-2852 in cell-lysate and (I) the effect of BI-2852 on purified KRAS. Cartoons are colored according to the color scale provided.

Considering that we were able to identify 94 individual peptide analytes for KRAS we continued using this method to acquire HDX measurements. HDX was initiated with a 9-fold dilution of the KRAS-containing cell lysate into labeling buffer (50 mM HEPES, 150 mM NaCl, pH 7.8, 99% D_2_O). Exchange reactions were quenched at specific timepoints with a 1:1 (v/v) addition of quench buffer (5 M guanidine hydrochloride, 0.8% formic acid, 2.5 % *n*-dodecyl-β-D-maltoside (DDM), 0.8 M Tris(2-carboxyethyl)phosphine (TCEP), and 1.5 mg *Aspergillus* protease, adjusted to pH 2.3). Quenched samples were injected immediately into the LC-MS system for online proteolytic digestion using an in-house packed pepsin column followed by RP-UPLC separation of resultant peptides and MS analyses.

HDX-MS experiments with KRAS in-lysate were performed both in the presence and absence of BI-2852, a small-molecule KRAS inhibitor (40). Prior to deuterium labeling, equilibration of BI-2852 binding was achieved with the addition of a 2-fold molar excess of the inhibitor (relative to the estimated concentration of KRAS in the lysate) to the cell lysate for 30 min. Direct comparison of HDX profiles for KRAS with and without BI-2852 revealed localized changes to deuterium incorporation for both the in-lyaste and purified KRAS (Figure 2F, H). Comparison of the BI-2852-induced differential HDX profiles between the two matrix conditions revealed comparable conformational changes in KRAS. Specifically, the differential HDX profile of BI-2852-bound KRAS measured directly in-lysate closely resembled that obtained using purified KRAS (Figure 2G, I). Importantly, BI-2852 induced decreased deuterium incorporation (although on different time-scales) in peptide regions that encompass switch II and the α4 helix in both purified and in-lysate KRAS; key aspects of the ligand-induced conformational response to inhibitor binding are retained in the cellular environment. Importantly, in-lysate KRAS displayed an additional region of HDX difference. There was increased deuterium incorporation within the N-terminus, nucleotide-binding region and the C-terminal region. These additional regions of difference in HDX suggest that, although the overall dynamic response of KRAS to BI-2852 is broadly conserved between purified and lysate conditions, the matrix components of the cellular environment introduce additional effects on backbone conformation that are not captured in the purified system. The chance that transient or stable interactions with cellular metabolites, soluble proteins, etc. is not zero. We are currently exploring which cellular component is responsible for these differences. Regardless, these results demonstrate that lysate-based HDX-MS can detect small-molecule-induced structural perturbations directly in a complex cellular milieu, providing an opportunity to examine how therapeutic ligands engage and remodel their protein targets under increasingly physiologically relevant conditions. Here we have used the lysate from an *E.coli* expression host as a proof-of-concept system, but we have data suggesting that this protocol will work with other expression host systems such as *Shewanella oneidensis MR-1* (data not shown).

### Is back-exchange a problem in a 20-minute gradient?

Greater sample complexity can compromise resolution through peptide co-elution, spectral complexity, and ion suppression necessitating a longer LC gradient for improved peptide separation. We therefore chose a 20-minute gradient for the crHDX workflow as described above. However, extending the chromatographic separation also increases the time that deuterated peptides are exposed to H₂O-containing mobile phases and could consequently increase deuterium back-exchange. To determine whether the extended LC gradient substantially affected the measured deuterium levels, purified KRAS was analyzed using a 5-35% acetonitrile:water (0.1% formic acid) gradient of 3 different lengths (6, 10, and 20 min) (Figure 3A). For a condensed visualization of the overall deuterium back-exchange behavior during three different LC gradient, KRAS peptides were grouped into four sequence regions encompassing residues 1-40, 41-80, 81-120, and 121-170. The relative percent deuterium uptake of peptides within each region was averaged at each labeling time point and compared (See also Supplemental excel data file). We observed minor differences among the three gradient conditions, indicating that extending the separation to 20 min did not substantially alter the measured deuterium levels under the conditions employed (Figure 3B, Supplemental excel data file). Based on these observations, the suitability of the 20-min gradient was confirmed for analyses, providing both improved separations for the more complex samples while maintaining acceptable deuterium retention. For MS acquisition, cIMS parameters were selected and optimized based on previously reported cIMS workflows for HDX analysis (22, 41), and the details are provided below (see Methods).

**Figure 3.**
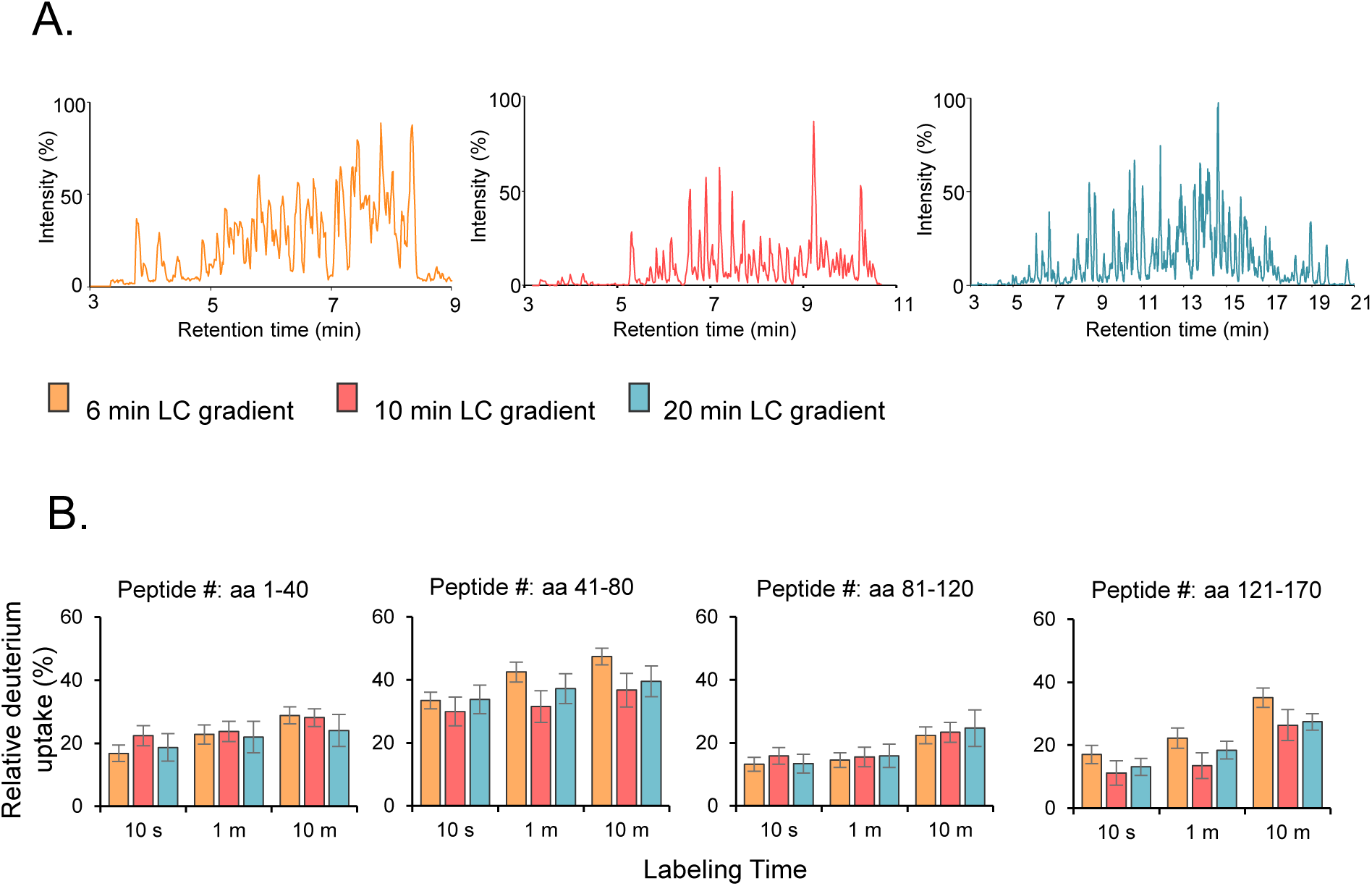
Optimization of the LC gradient for crHDX analysis. **(A)** (A) Representative LC-MS chromatograms of purified KRAS peptides separated using a 5–35% acetonitrile:water gradient over three different gradient durations (6, 10, and 20 minutes). **(B)** Purified KRAS was analyzed using increasing LC gradients to evaluate the effect of extended chromatographic separation on measured deuterium incorporation. KRAS peptides were grouped into four sequence regions encompassing residues 1–40, 41–80, 81–120, and 121–170. The relative percent deuterium incorporation of the peptides within each region was averaged at each labeling time point and compared across the three LC gradients. The error bars represent calculated standard error of mean (SEM).

### Development and optimization of crHDX in complex cellular environments: in-cells

Confident that we were able to perform successful HDX-MS experiments in-lysates without isolation of the target protein away from the matrix, we sought to develop a complementary in-cell HDX labeling approach. To both develop and evaluate this, we again selected *E. coli* BL21(DE3) cells overexpressing His₆-tagged KRAS as our model system. Choosing a common expression host system found in many research laboratories was intentional and we believe that our protocol will work with other common laboratory expression host systems as well. The workflow was organized into four sequential steps prior to the fifth LC-MS analysis step (Figure 4A) and we present our detailed protocol below.

**Figure 4.**
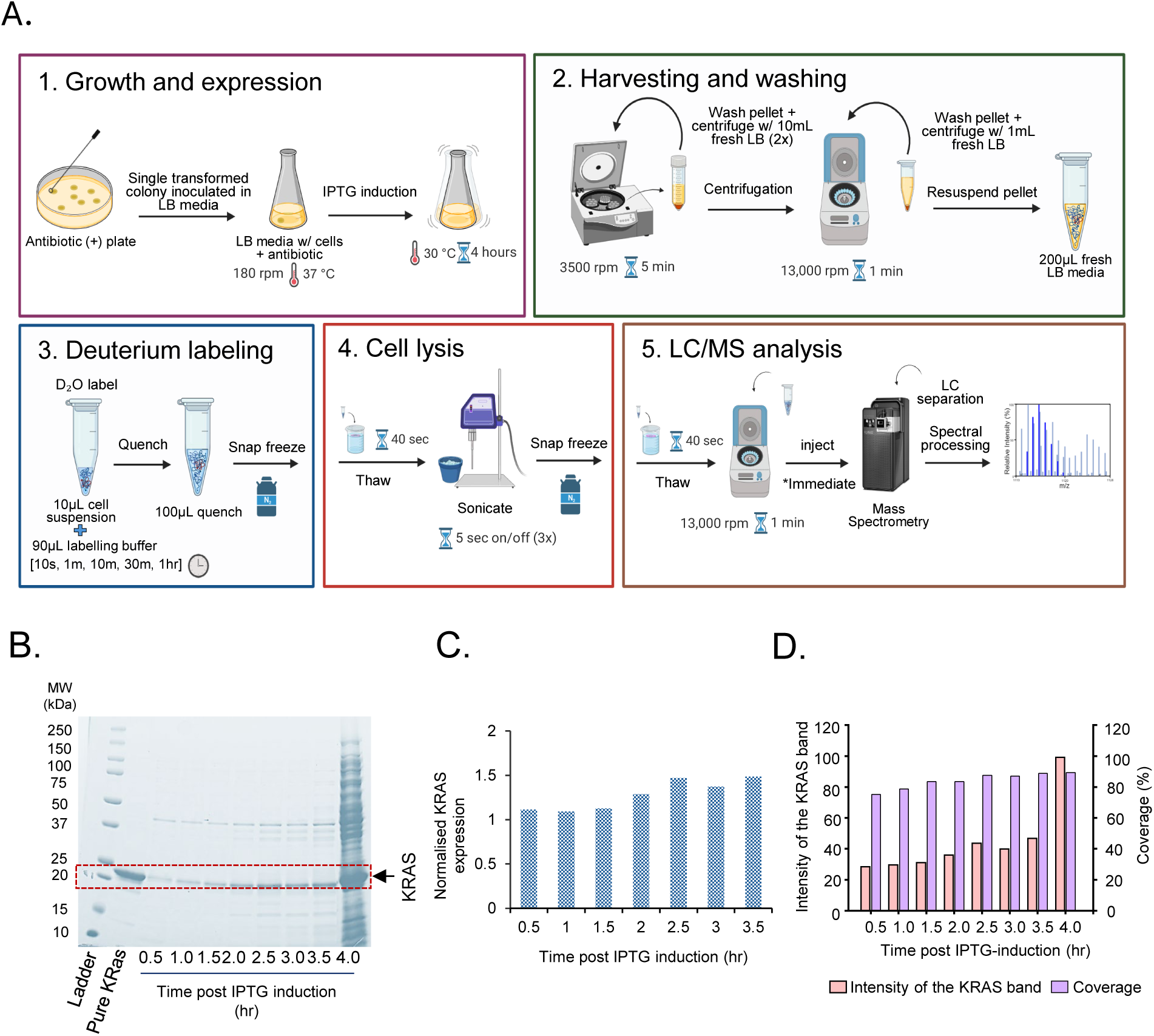
Stepwise development and optimization of the in-cell crHDX workflow. **(A)** Schematic representation of the five-step in-cell crHDX workflow developed for monitoring the conformational dynamics of an overexpressed target protein directly in intact *E. coli* cells. *Step 1: Growth and expression (magenta box)*, His₆-tagged KRAS (1–169) was overexpressed in *E. coli* BL21(DE3) cells following IPTG induction. *Step 2: Harvesting and washing (dark green box)*, cells were harvested by centrifugation and gently washed to remove residual culture medium while minimizing disruption of cell integrity. *Step 3: Deuterium labelling (blue box)*, the optimized amount of cell suspension was subjected to deuterium labeling by dilution into D₂O-containing buffer for defined exchange time points, followed by quenching and immediate snap-freezing. *Step 4: Cell Lysis (red box)*, labeled cells were rapidly thawed and disrupted by controlled sonication on ice, followed by immediate snap-freezing to minimize deuterium back-exchange. *Step 5: LC/MS analysis (brown box)*, lysed samples were rapidly thawed, clarified by centrifugation, diluted under acidic conditions, and immediately subjected to LC-MS analysis. **(B)** SDS-PAGE analysis of *E. coli* cell pellets collected at different times following IPTG induction, demonstrating the progressive increase in KRAS expression. The highlighted region (red box) indicates the band corresponding to overexpressed KRAS. **(C)** Densitometric quantification of KRAS expression from SDS-PAGE analysis at different post-induction time points. KRAS band intensity was quantified and normalized to the combined intensity of the remaining protein bands within the corresponding sample lane to assess relative KRAS expression levels over time. **(D)** Effect of post-induction expression time on KRAS peptide identification and HDX-MS analysis using the crHDX workflow. The bar graphs show the abundance of KRAS at different post-induction expression time from the SDS-PAGE densitometry analysis (light orange) and its sequence coverage (purple) obtained from HDX-MS analysis. For calculation see Supplemental excel, tab Figure 4.

### The in-cell crHDX method (steps 1-4)

In the first and second step, the KRAS construct, cloned into the pET-28a(+) expression vector residues 1-169, was overexpressed in *E. coli* BL21(DE3) cells and the resultant cells were prepared for deuterium labeling. First, bacterial starter cultures were grown overnight in LB medium at 37 °C containing 0.1 mg/mL kanamycin. After the overnight growth, the culture was diluted 50-fold into fresh medium and grown at 37 °C until an OD₆₀₀ of approximately 0.5 was reached. KRAS expression was then induced with addition of 0.25 mM IPTG, followed by incubation at 30°C for 4 h with shaking (Figure 4A, Step 1). In the second step, a 10 mL aliquot of the induced cell culture was centrifuged at 3,500 rpm for 10 min at 4°C. The resulting cell pellet was gently resuspended and washed with 5 mL of fresh LB medium. This washing procedure was repeated twice using the same protocol. Gentle handling was maintained throughout this step, as well as subsequent steps, to minimize unintended cell disruption and preserve cell integrity prior to deuterium labeling. Following the final wash, the cell pellet was resuspended in 1 mL of fresh LB medium and centrifuged at 13,000 rpm for 1 min. The resulting pellet was immediately and gently resuspended in 200 µL of fresh LB medium for subsequent HDX labeling (Figure 4A, Step 2). The above protocol might not be amenable to other cell types or POIs so the washing conditions and volumes will need to be further optimized depending on the host used, the cell culture density, and the experimental requirements.

In the third step, the amount of cell suspension that contains sufficient target protein for control sample MS detection and subsequent analyses of deuterium labeled peptides is defined using undeuterated control samples. We used total protein concentrations, estimated by Bradford assay using different volumes of the cell suspension, and SDS-PAGE analysis to assess the relative abundance of overexpressed KRAS and to determine the amount of cell suspension required to obtain sufficient KRAS for MS analysis. Based on these analyses, 10 µL of the concentrated cell suspension after 4 hours of induction, corresponding to approximately 15 µg of total protein, was selected for each HDX reaction to ensure > 90 % sequence coverage and redundancy for KRAS. From the SDS-PAGE analysis of the 4-hour induction sample the overexpressed KRAS was determined to be a major component of the protein expression profile [approximately 90%] (Figure 4B, Supplemental Excel; tab: Figure 4), suggesting that this volume would provide sufficient quantity of target protein for subsequent HDX-MS analyses. In conventional HDX-MS analyses, 10-20 picomoles of KRAS per injection are required; this is in line with the quantity estimated from the above. As such, for undeuterated control samples, 10 µL of cell suspension was diluted with 90 µL of H₂O buffer (20 mM HEPES, 150 mM NaCl, 1 mM DTT, and 5 mM MgCl₂, pH 7.8). For deuterated samples, 10 µL of the cell suspension was diluted with 90 µL of labeling buffer prepared in 99% D₂O (same buffer composition as H_2_O buffer above), resulting in a maximum D₂O content of 89.1% during the exchange reaction. After labelling for specific amounts of time, an equal volume, v/v (100 µL) of quench buffer (5M Guanidium Hydrochloride, 0.8 % Formic acid, 0.8 M TCEP, 1.5 mg of *Aspergillus* protease enzyme and 2.5 % DDM, pH 2.23) was added to quench HDX. Quenched samples were immediately snap-frozen and stored at −80°C until further processing (Figure 4A, Step 3).

In the fourth step, the frozen, quenched cell samples are thawed for 40 s in a 22°C water bath, briefly centrifuged, and immediately placed on ice. The cells are then subjected to sonication for a total of 15 s using alternating 5-s sonication and 5-s pause cycles. Sonication was performed on ice to maintain the sample at approximately 0°C minimizing further back-exchange during cell disruption. Immediately following sonication, the lysed samples were snap-frozen until further processing (Figure 4A, Step 4). In our hands, Step 4 takes approximately 1 minute. A recent study suggests that in this amount of time the increased level of back-exchange is negligible (42).

### The in-cell crHDX LC-MS analysis (step 5)

As the fifth and final step, we needed to confirm that our LC-MS protocol that we optimized for use with unfrozen cell lysate would be appropriate for this application. The frozen cell lysate prepared in step 4 was thawed for 40 s in a 22°C water bath followed immediately by centrifugation at 13,000 rpm for 1 min to remove both the insoluble cellular material and the material that precipitated as a result of the freeze-thaw cycle. Based on the amount of protein determined to be suitable for the LC-MS analysis during the initial optimization, 5 µL of the supernatant was used for each injection. This aliquot was then diluted with 20 µL of 0.1% formic acid in water to obtain a final sample volume of 25 µL, compatible with the 50 µL injection loop used in our LC-MS setup. The quenched sample prepared was then immediately injected into the LC system for further online digestion, LC separation and MS analyses (Figure 4A, Step 5) as before. All together, these steps modified from the traditional HDX-MS experiment enabled successful recovery and LC-MS detection of KRAS-derived peptides following deuterium labeling directly in-cells, establishing the experimental workflow for subsequent in-cell crHDX analysis.

### Is 4-hours of overexpression post IPTG-induction required?

We also investigated how the abundance of the target protein influenced its detection by our crHDX workflow. Previous in-cell HDX-MS labeling relied on the natural abundance of the POI over that of the other host cell proteins (32). For initial method development, we chose to use the 4-hour induction time-point as we were confident that there would be enough target protein for analysis. However, we were interested to determine if we could use earlier induction time-points for analyses thereby minimizing time-to-analyses, as well as getting an understanding of the limits of the method; not all POIs and their variants express to the same levels. To generate progressively different levels of KRAS expression, cells were collected at 30-min intervals following IPTG induction for up to 4 h (Figure 4B). As expected, SDS-PAGE analysis showed a progressive increase in the abundance of KRAS over the induction period (Figure 4B). We also determined the intensity of KRAS expression against host-cell proteins for each IPTG-induction time point as in Figure 4B. This example illustrates that the induction time did not appear to have a significant difference in the abundance of KRAS when compared to the host-cell proteins (Figure 4C). Each induction time point was subsequently processed using our crHDX workflow above to determine whether sufficient KRAS peptides could be recovered during MS analysis (Figure 4D). Remarkably, KRAS could be detected with substantial sequence coverage even at the earliest induction time point. After only 30 min of IPTG induction, the expression of KRAS was sufficient to identify peptides with 75.1 % sequence coverage, demonstrating that very high levels of target-protein overexpression were not required for effective peptide detection under the conditions tested (Figure 4D). However, increasing the induction period beyond 2.5 hours resulted in robust KRAS production with sequence coverage exceeding 80%, indicating consistently broad peptide representation of the KRAS sequence as its cellular abundance increased (Figure 4C, 4D). Most importantly, our workflow also remained effective at all induction time points despite the increasingly complex and concentrated cellular protein background. After 4 h of induction, when SDS-PAGE showed a highly complex and concentrated protein profile in addition to strongly overexpressed KRAS, the crHDX analysis provided 89.4% KRAS sequence coverage with a peptide redundancy of 5.57. Thus, increasing cellular sample complexity did not prevent extensive recovery and identification of KRAS-derived peptides when the target protein was present at sufficient abundance. Together, these results demonstrate that our crHDX workflow operates across a range of target-protein expression levels. The ability to achieve nearly 80% sequence coverage at time points, prior to 4h post IPTG induction indicates that the method does not always require maximal overexpression of the target protein, while the high sequence coverage and redundancy observed after prolonged induction demonstrate that the workflow remains effective within a highly complex cellular protein background. These findings further define the practical operating range of the method and suggest that target abundance, rather than isolation to near homogeneity, may be a key determinant of successful crHDX analysis.

### crHDX for Monitoring Protein Dynamics in-Cells

Previous studies have demonstrated that D₂O can readily enter cells, supporting the feasibility of deuterium labeling within an intact cellular environment (31, 32, 43). We therefore evaluated whether the developed protocol could reliably capture KRAS structural dynamics when deuterium labeling was performed in intact bacterial cells prior to cell disruption. For this we compared the HDX-MS profiles of KRAS across three distinct experimental conditions: (i) KRAS followed by cell disruption after deuterium labeling (in-cell labelling) (ii) KRAS analyzed directly in the unfractionated cell lysate (in-lysate) and (iii) conventionally purified KRAS. As a first step in comparison, we compared the chromatographic separation (Figure 5B) and the proteolytic digestion profiles (Figure 5C) obtained for KRAS across the three experimental conditions. The comparison of the BPI chromatograms of peptide separations across the three experimental conditions revealed distinct differences as sample complexity increased. Purified KRAS displayed a relatively well-resolved chromatographic profile, with KRAS-derived peptides distributed across the 20-min LC gradient (peptides eluted between 7.0 - 17.0 minutes) with average peak widths measured as on an average of 0.016 minutes (FWHM). In contrast, KRAS analyzed in the unfractionated cell lysate showed the increasingly crowded chromatographic profile as this sample includes the peptides derived from numerous soluble host-cell proteins. The complexity increased again in samples obtained from in-cell crHDX. However, KRAS-derived peptides eluted similarly in a range between 7.0-17.0 minutes and exhibited an average peak width of 0.018 minutes and 0.010 minutes (FWHM) for in-lysate and for in-cell, respectively (Supplemental excel, tab; Peak Width) despite a greater number of peptide signals distributed across the linear separation. Interestingly, in addition to, and perhaps as a result of, the increased chromatographic crowding the distribution of some of the KRAS-derived peptides shifted towards earlier retention times, while maintaining IM drift time, in the more complex lysate and in-cell samples compared with that of the purified KRAS (Figure 5B, Supplemental excel, tab; Peak Width). The grey boxes in the in-lysate and in-cell chromatograms highlight the elution regions for a majority of KRAS peptides that were routinely identified across both these samples and purified KRAS. We believe that the altered chromatographic behavior is due to the analyte-matrix-dependent effects associated with the increased complexity of these samples; co-eluting peptides from host-cell proteins, competition for interactions with the stationary phase, differences in peptide loading, and the presence of additional cellular components could potentially influence peptide retention and contribute to the observed shift in the KRAS chromatographic profile.

**Figure 5.**
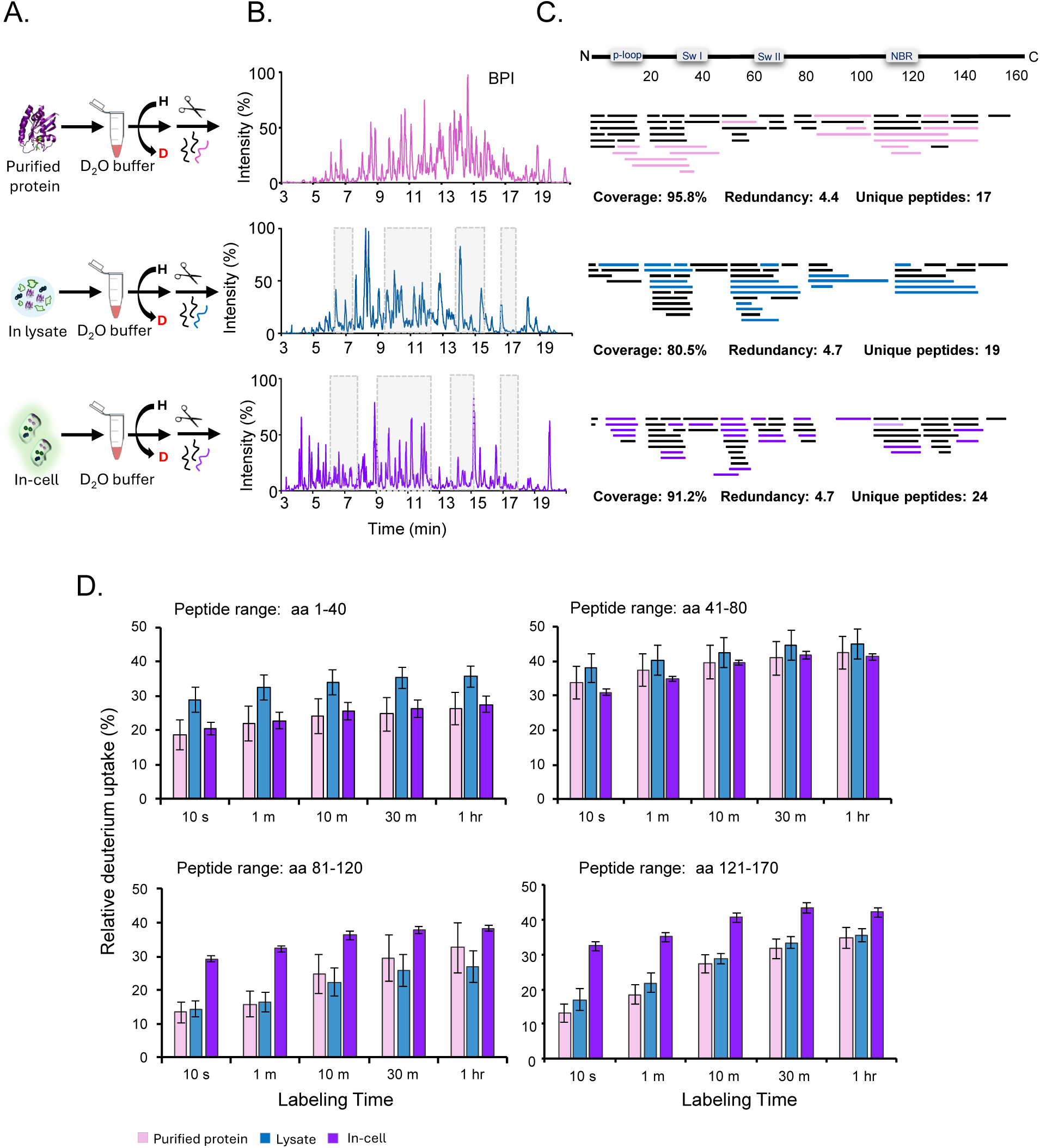
Validation of the in-cell crHDX workflow by comparison of KRAS across purified, lysate, and intact-cell environments. **(A)** Schematic representation of the three experimental conditions used for comparative HDX-MS analysis: purified KRAS in buffer, KRAS in cell lysate, and KRAS labeled directly within intact *B. coli* cells prior to cell disruption. **(B)** Representative LC chromatograms obtained for KRAS under the three experimental conditions. Highlighted regions indicate representative retention-time windows shows the KRAS peptide elution. **(C)** Peptide coverage maps of KRAS (excluding the His_6_ tag and the linker) obtained from purified protein, unfractionated cell lysate, and intact-cell samples. Individual horizontal black bars represent KRAS peptides found in all three experimental conditions. The Pink-, blue- and purple-colored bars correspond to the unique peptides for the three different biological conditions respectively. Major structural regions, including the p-loop, switch I (sw I), switch II (sw II), and nucleotide-binding region (NBR), are indicated. **(D)** Comparison of the time-dependent deuterium uptake of KRAS across the three experimental environments. For simplified visualization, identified KRAS peptides were grouped into four sequence regions encompassing residues 1-40, 41-80, 81-120, and 121-170. Relative percent deuterium uptake values for peptides within each region were averaged at each labeling time point and compared among purified KRAS, lysate KRAS, and intact-cell KRAS.

We next examined whether these differences in chromatographic complexity were also reflected in the peptide populations recovered for KRAS under each condition. Comparison of the peptides identified revealed that 20 peptides were common across all three conditions (Figure 5C, black colored bars) spanning 68.6 % of the KRAS sequence. Each condition also produced a number of unique peptides: purified KRAS: 17; KRAS from in-lysate: 19; and KRAS from in-cell: 24 peptides (Figure 5C and Supplemental excel tab: Figure 5). These differences likely reflect the distinct sample environments in which the HDX experiment was performed, ranging from purified protein in a defined buffer to in-lysate and, ultimately, in-cells. The surrounding sample matrix can influence peptide generation and detection; these include proteolytic digestion efficiency, peptide recovery, chromatographic separation, ionization efficiency, spectral complexity post deuterium incorporation, and ion suppression. Despite the substantially greater complexity of the in-cell crHDX samples, the KRAS peptides obtained provided sequence coverage and peptide redundancy that was comparable to those obtained from KRAS both in the in-lysate and also purified KRAS (96, 81, and 91% coverage and 4.4, 4.7, and 4.7 for purified, in-lysate, and in-cell, respectively). Sufficient KRAS sequence coverage was maintained in all cases.

We next evaluated whether time-dependent deuterium exchange could be reliably captured using the crHDX workflow in-cell. To confirm this, deuterium incorporation into KRAS was monitored across different labeling time points for each experimental condition and in replication (Figure 5D, all HDX data are contained in Supplemental data excel tab:Figure 5). As above, we chose to visualize the comparison through a binned relative percent deuterium exchange behavior. As in Figure 3B, KRAS peptides were combined into four sequence regions encompassing residues 1-40, 41-80, 81-120, and 121-170 as they covered the full sequence of KRAS. The relative percent deuterium incorporation of peptides within each region was averaged at each labeling time point and compared across KRAS labeled in in-cells, in-lysate, and purified protein (Figure 5D). The region encompassing residues 41-80 showed broadly comparable HDX across all three conditions. In contrast, KRAS labeled within intact cells displayed increased HDX within residues 81-120 and 121-170 relative to KRAS analyzed in both the lysate and purified states. The N-terminal region encompassing residues 1-40 showed mildly decreased deuterium incorporation for in-cell KRAS compared with in-lysate KRAS, while being more comparable to purified KRAS (Figure 5D). Importantly, despite the region-specific differences in HDX, the overall exchange behavior of KRAS remained comparable across the three experimental states. Together, these results demonstrate that the workflow that we developed can reliably capture protein conformational dynamics within increasingly complex cellular and analytical environments. We are currently exploring the cellular components that might be affecting the deuterium incorporation of KRAS under each of these conditions.

### Extending the crHDX workflow to investigate protein-protein interactions in-cells

As the crHDX workflow could reliably capture the conformational dynamics of an overexpressed target protein, we next asked whether the approach could be extended to investigate protein-protein interactions directly within a cellular environment without being able to control for exact binding conditions; *in vitro* HDX-MS experiments conventionally maximize ligand occupancy. To address this, we explored a more complex system in which two proteins that are known to interact were co-expressed within the same bacterial cells. Here, we selected KRAS and the catalytic domain of Son of Sevenless (SOS_cat_) as our model protein-protein interaction system. SOS is a guanine nucleotide exchange factor (GEF) that interacts with KRAS and promotes nucleotide exchange by facilitating the release of GDP, thereby enabling subsequent GTP binding and activation of KRAS (44). In our recent HDX-MS study, we have shown that the KRAS interaction with SOS_cat_ induces distinct changes in backbone conformational dynamics for both proteins (34). This well-characterized conformational response provides an appropriate benchmark to test whether our crHDX workflow can detect protein-protein interaction-induced conformational changes in KRAS directly within the cellular environment. For this, constructs encoding both His₆-tagged KRAS and His₆-tagged SOS_cat_ were co-transformed into *E. coli* cells and co-expressed, allowing the two proteins to coexist within the same cellular environment prior to deuterium labeling. Individual colonies were screened by SDS-PAGE to confirm the simultaneous expression of both KRAS and SOS_cat_ (Figure 6A). A colony showing robust expression of both proteins was selected and subsequently grown into a larger-scale culture. The cells co-expressing KRAS and SOS_cat_ were then subjected to the crHDX workflow (Figure 4A, Figure 6). In the present study, *E. coli* was used as the host-cell as a proof-of-concept system to establish and validate our crHDX workflow in detecting protein: protein interactions, but we believe that additional expression hosts can be used as well.

**Figure 6.**
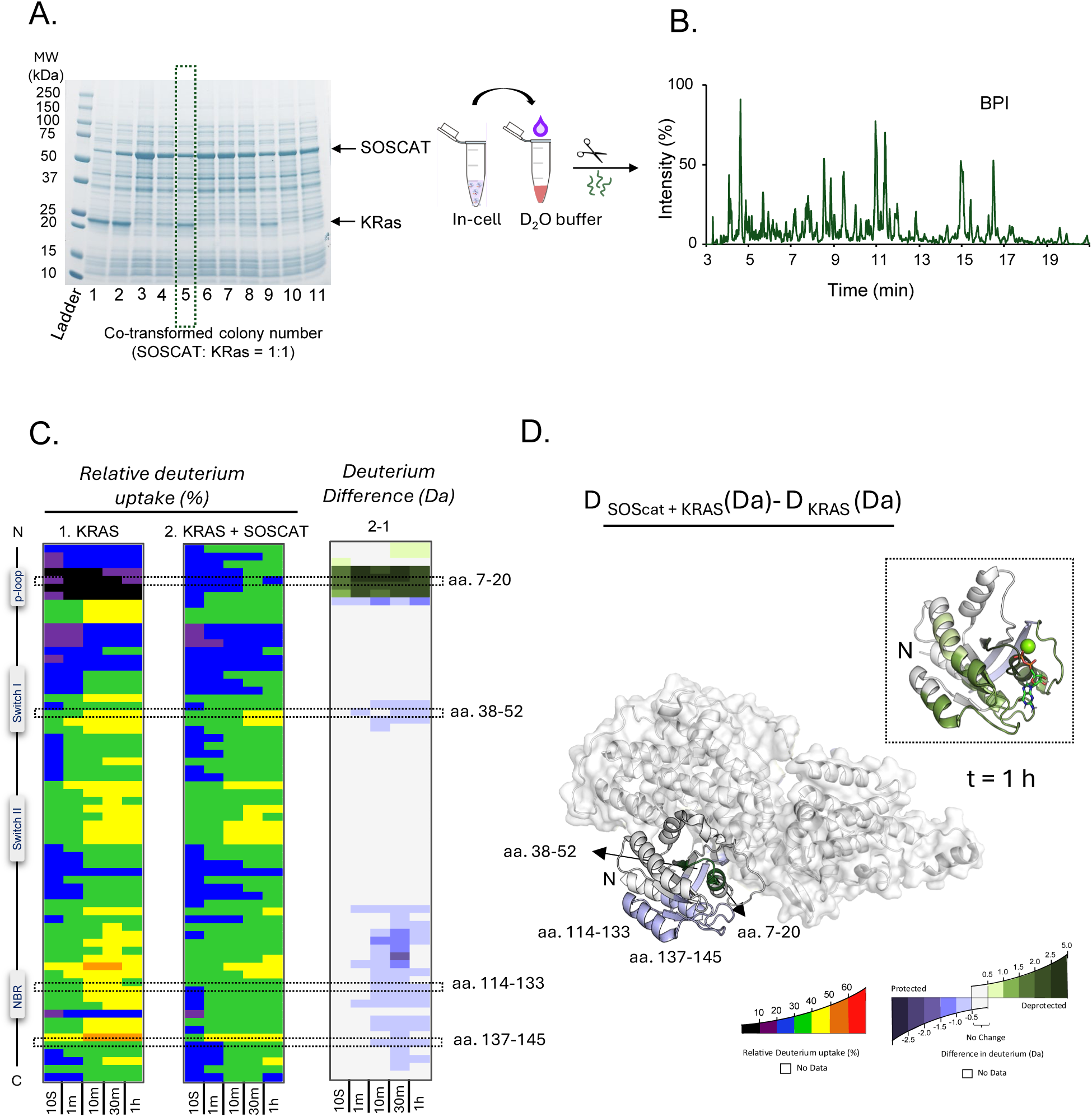
In-cell crHDX captures SOS_cat_-induced conformational changes in KRAS. **(A)** SDS-PAGE analysis of E. coli colonies co-transformed with constructs encoding His₆-KRAS and SOS_cat_. Colonies were screened for simultaneous expression of both proteins, and a clone exhibiting robust co-expression of both POIs was selected for subsequent in-cell crHDX analysis. Schematic representation of the in-cell protein-protein interaction experiment. KRAS and SOS_cat_ were co-expressed within the same cellular environment, followed by deuterium labeling of in-cells and processing using the optimized crHDX workflow. **(B)** Representative LC-MS profile obtained from cells co-expressing KRAS and SOS_cat_. **(C)** Peptide-level HDX-MS analysis comparing KRAS expressed alone versus KRAS co-expressed with SOS_cat_. Deuterium uptake profiles and differential HDX maps illustrate SOS_cat_-induced changes across KRAS, with the p-loop, switch I, switch II, and nucleotide-binding region (NBR) indicated. **(D)** Differential HDX changes mapped onto the KRAS-SOS_cat_ structural model (PDB: 7KFZ). Regions exhibiting increased or decreased deuterium uptake upon SOS_cat_ co-expression are represented according to the indicated color scales. The inset shows the differential deuterium uptake mapped onto KRAS from our previous *in vitro* KRAS-SOS_cat_ HDX-MS study (34), allowing direct comparison of SOS_cat_-induced conformational changes observed under *in-cell* and purified *in vitro* conditions.

To assess the conformational effects of SOS_cat_ on KRAS backbone dynamics in-cells, the deuterium incorporation profile of KRAS in cells co-expressing both KRAS and SOS_cat_ was compared to that of cells expressing only KRAS (Figure 6C). Co-expression with SOS_cat_ resulted in increased deuterium incorporation within the N-terminal region of KRAS, together with decreased deuterium incorporation within the switch I region. Importantly, these changes in HDX were consistent with the changes in HDX observed for the KRAS-SOS_cat_ interaction using purified proteins as described in our previous *in vitro* study (34) (Figure 6D). The ability to reproduce these previously characterized SOS_cat_-induced changes provided further evidence that the crHDX workflow can capture protein-protein interaction-dependent alterations in KRAS conformational dynamics. Interestingly, the in-cell measurements also revealed decreased deuterium incorporation within the C-terminal region of KRAS upon SOS_cat_ co-expression, a response that is in direct contrast to that which was observed in our previous *in vitro* study (Figure 6C, D). This difference suggests that the conformational response of KRAS to SOS_cat_ may be influenced by the complexity of the intracellular environment. Cellular crowding, endogenous molecular interactions, differences in nucleotide occupancy, or other components present within the cell could potentially modulate the structural response of this region during the KRAS-SOS_cat_ interaction. Most importantly, while the in-cell measurements reproduced key features of the previously characterized *in vitro* interaction, they also revealed region-specific differences that may not be captured when the proteins are examined in isolation. We are currently working to identify the cellular components that could be contributing to this altered HDX response.

### Expanding crHDX beyond conventional protein purification requirements

We next explored the application of our in-cell HDX workflow to proteins for which conventional purification may be challenging or potentially undesirable. Affinity tags, for one, can be problematic for some proteins. Although affinity tags substantially facilitate protein purification, their presence may not be suitable for every protein (45, 46) and, in some cases, could potentially influence protein conformational dynamics (Figure 7A). Alternatively, purification of an untagged protein often requires multiple chromatographic and or precipitation steps (all depending on the physiochemical properties of the target protein) to achieve the level of purity that is typically required for conventional HDX-MS. Therefore, we tested whether our in-cell workflow could be applied directly to an overexpressed, untagged target protein. Similar to the above, here we chose to express untagged KRAS in *E. coli* cells and subjected the cells to the optimized crHDX workflow (Figure 4A). Despite the absence of an affinity tag, KRAS peptides were readily detected, providing substantial sequence coverage suitable for HDX-MS analyses (Figure 7B, Supplemental excel tab; Figure 7). Subsequently, we performed deuterium labeling over multiple exchange time points and successfully monitored time-dependent deuterium incorporation into untagged KRAS within the *E. coli* cells. These results demonstrate that the workflow is not enabled by the presence of a His_6_ affinity tag and thus provides an alternative strategy for investigating conformational dynamics for those proteins for which tagging is undesirable or where purification of the untagged protein would otherwise require extensive chromatographic processing.

**Figure 7.**
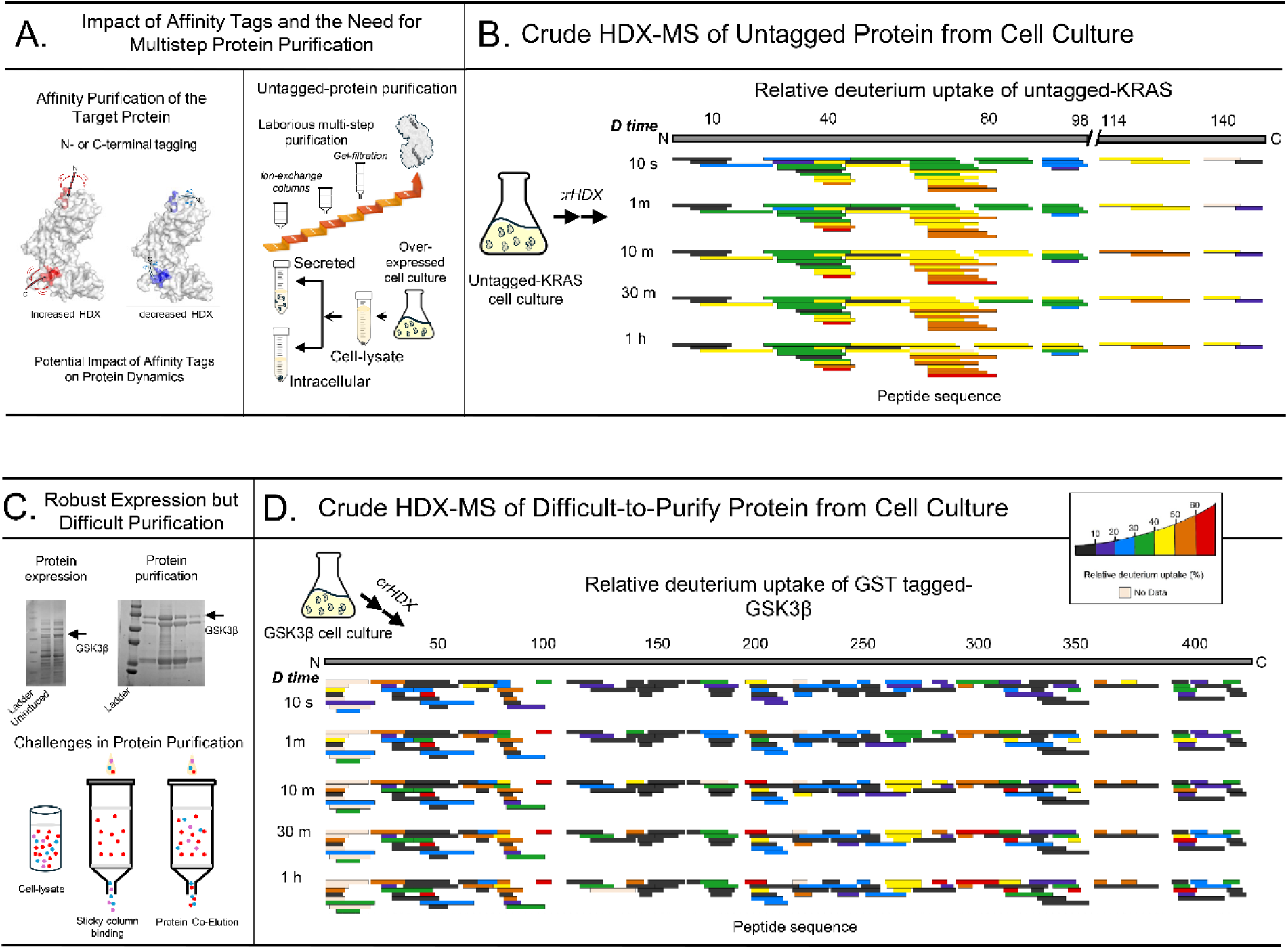
Application of CrHDX to Untagged and Difficult-to-Purify Proteins: **(A)** Schematic illustrating potential limitations associated with affinity tagging and conventional purification. Introduction of an affinity tag may influence protein structure or molecular interactions, whereas purification of untagged proteins can require multiple chromatographic steps. **(B)** Application of crHDX to untagged KRAS. Untagged KRAS was expressed in *E. coli* and subjected to the crHDX workflow. Peptide coverage maps across different deuterium-labeling time points demonstrate broad sequence representation of untagged KRAS, supporting the applicability of the method to proteins for which affinity tagging may be undesirable. Individual horizontal bars represent peptides detected at the indicated labeling time points. **(C)** Application of crHDX to a purification-challenging protein GSK3β. SDS-PAGE analysis illustrates robust expression of GSK3β in *E. coli* despite difficulties encountered during conventional purification due to co-elution of other protein. **(D)** Peptide coverage maps of GSK3β obtained across the HDX labeling time course. Individual horizontal bars represent peptides detected at the indicated labeling time points. Extensive peptide representation across the GSK3β sequence demonstrates that crHDX can recover target-derived peptides even for a protein that is difficult to obtain in a highly purified form.

Another practical challenge in conventional HDX-MS comes with proteins that are efficiently overexpressed but remain difficult or impossible to purify to usable or measurable quantities. Such proteins may exhibit poor compatibility with affinity, or other, column purification or nonspecific column interactions resulting in co-elution with other proteins (47, 48), aggregation, or loss of significant soluble protein during purification (49, 50). In these cases, obtaining sufficient quantities of highly pure and structurally intact protein can become a major hurdle in person-time and cost limiting their likelihood for structural analyses. To evaluate whether the crHDX workflow could overcome this limitation, we selected glycogen synthase kinase 3 beta (GSK3β) as a purification-challenging model protein. Although GSK3β was efficiently overexpressed in *E. coli*, purification to homogeneity was challenging as the target consistently co-eluted with other highly abundant proteins of similar apparent molecular weight (Figure 7C). We therefore applied our crHDX workflow directly to cells overexpressing GSK3β, bypassing the requirement of chromatographic purification prior to deuterium labeling (Figure 7D). Considering that the in-cell, and untagged HDX-MS data for KRAS align well with previously acquired HDX data, we are confident that the data we have acquired for GSK3β are representative of its backbone dynamics within the complex environment of the expression host.

## Conclusions

In the present study, we have developed a crude HDX-MS workflow that extends conventional HDX-MS away from purified proteins towards complex cellular environments. Using KRAS, a well-characterized model system in our laboratory (34–37), we demonstrated that an overexpressed target protein can be analyzed directly from within complex protein mixtures with sufficient sequence coverage and peptide redundancy for comparative HDX-MS analysis. Importantly, deuterium labeling was performed directly in intact *E. coli* cells, followed by controlled cell disruption, proteolytic digestion, and LC-MS analysis, enabling time-dependent protein conformational dynamics to be measured within a cellular environment. Comparison of KRAS under purified, in-lysate, and in-cell conditions revealed broadly comparable HDX behavior across several peptide regions, while distinct region-specific differences emerged in the in-cell experiment. These differences suggest that the conformational ensemble measured in-cells differs from that observed after either or, or both, cell disruption or purification; this might reflect the combined influence of macromolecular crowding, metabolites, ions, nucleotide state, and endogenous molecular interactions. Importantly, the applicability of our crHDX methods extended beyond monitoring the dynamics of an individual protein. Co-expression of KRAS and SOS_cat_ demonstrated that protein-protein interaction-dependent conformational changes can be detected directly in-cells. Several SOS_cat_ induced changes in KRAS were consistent with those observed previously using purified proteins (34), providing an important benchmark for the workflow, while additional region-specific differences were observed under in-cell conditions. Together, these findings show that crHDX can bridge conventional purified-protein HDX-MS and measurements performed in more biologically complex environments, enabling the direct comparison of protein dynamics and molecular interactions across different levels of cellular complexity. The workflow also addresses practical limitations associated with purification-dependent HDX-MS. Successful analysis of untagged KRAS demonstrated that affinity tagging is not inherently required when the target protein is present at sufficient abundance, expanding the approach to systems in which a tag may perturb protein structure, function, or molecular interactions. While KRAS can be purified as tagged or untagged, our HDX application to in-cell GSK3β demonstrated the feasibility of examining proteins that are readily expressed but difficult to purify to near homogeneity because of co-elution, unfavorable chromatographic behavior, aggregation, or protein loss during purification. Collectively, these results establish crHDX as a complementary strategy to conventional HDX-MS; there is no longer the absolute requirement for extensive purification while retaining the ability to obtain protein-specific, time-resolved information on conformational dynamics and molecular interactions.

While *E. coli* was used here as a proof-of-concept system, the broader opportunity for crHDX lies in extending the approach to other cellular systems and increasingly diverse classes of proteins. Future studies could expand beyond overexpressed model proteins to examine endogenous host-cell proteins, including enzymes, signaling proteins, structural proteins, and components of larger macromolecular assemblies. In fact, despite only analyzing the HDX data for the POI in each case, there are a plethora of data still left to mine, and in our case most specifically from the protein production machinery. Such applications could enable thew study of conformational dynamics in the context of cellular processes rather than exclusively at the level of an isolated protein. Particularly exciting would be the ability to simultaneously monitor multiple proteins participating in the same pathway or molecular complex, providing insight into how conformational changes and protein interactions are coordinated during a cellular response. Achieving this level of analysis will require further improvements in sensitivity, peptide identification, chromatographic and ion-mobility separations, and quantitative approaches capable of resolving multiple proteins with high sequence coverage within highly complex cellular backgrounds.

Nevertheless, crHDX measurements described here do represent ensemble-averaged conformational behavior across a population of cells. Individual cells can differ in protein abundance, metabolic state, interaction networks, and response to environmental or biochemical perturbations; consequently, conformational states present only within specific cellular subpopulations may be obscured by population averaging. Whether protein structural dynamics exhibit measurable cell-to-cell heterogeneity therefore represents an important future question. Although single-cell HDX-MS remains beyond the sensitivity of our current workflow, continued advances in mass-spectrometric sensitivity, low-volume sample handling, and single-cell proteomics may ultimately make it possible to interrogate protein dynamics in progressively smaller cell populations and, eventually, at the single-cell level. Together, these developments could extend HDX-MS from the characterization of individual purified proteins toward understanding how protein conformational dynamics and molecular interactions are organized and coordinated within cellular systems.

### Summary

This study establishes crHDX as a complementary HDX-MS workflow for investigating protein conformational dynamics directly in complex cellular environments while minimizing extensive protein purification. The approach enables analysis of intact-cell protein dynamics, protein-protein interactions, untagged proteins, and purification-challenging targets while maintaining useful sequence coverage and peptide redundancy. However, current limitations include dependence on sufficient target-protein abundance, increased chromatographic complexity, ion suppression, variable peptide recovery, and potential back-exchange during sample processing. Further improvements in MS sensitivity and separation could expand its application to endogenous and disease-associated upregulated proteins.

## Methods and Materials

### Protein expression and purification

The gene encoding human WT-KRAS (residues 1–169, Uniprot P01116), WT-SOS cat (558–1049, Uniprot Q07889), GSK3β (Uniprot ID: P49841) and Glucokinase (Uniprot ID: P35557) was synthesized and inserted into a plasmid to introduce either an N-terminal His tag (for KRAS, SOSCAT and Glucokinase) or a GST tag (for GSK3β) for affinity purification (GenScript). Similarly, WT KRAS (1–169) containing no tag was also cloned as described above. Constructs cloned in pET 28a(+) for KRAS and SOS_cat_ and pGEX3 for GSK3β were expressed using Escherichia coli BL21 (DE3) cells. Starter cultures were grown overnight in LB media containing 0.1 mg/mL kanamycin and ampicillin respectively. Overnight cultures were diluted 50-fold and grown at 37°C to A _600_ 0.5, then induced with 0.25 mM Isopropyl-D-1-thiogalactopyranoside (IPTG) for over-expression of protein at 30°C for 4 h with shaking. At this stage, a portion of the IPTG-induced bacterial cells in LB media was harvested to carry out the crude HDX protocol. The remainder of the bacterial cells were pelleted using centrifugation at 7000 RPM for 15 min. The collected pellet was resuspended in lysis buffer containing 20 mM Tris HCl, pH 7.6, 250 mM NaCl, 20 mM imidazole, and 2.5 mM PMSF and then subjected to cell lysis on ice. The lysate was then clarified using high-speed centrifugation at 19,000 RPM for 1 h. An aliquot of the supernatant was set side, followed by total protein estimation using the Bradford assay to carry out the HDX analysis. The remaining supernatant was further purified using Nickel affinity chromatography following standard protocols. The protein was eluted with 250 mM imidazole-containing buffer. The eluted protein was then subjected to size-exclusion chromatography using Superdex 75 10/300GL column (GE Healthcare Life sciences). Fractions containing pure protein were collected, concentrated, and the concentration was confirmed using the Bradford assay.

### Densitometric analysis

Protein expression levels were quantified by the densitometric analysis of SDS-PAGE band intensities using ImageJ software (version 1.xx; National Institutes of Health, Bethesda, MD, USA). After electrophoresis and staining, gels were imaged under uniform illumination and saved as high-resolution, uncompressed JPEG files to preserve band intensity information. Images were opened in ImageJ and converted to 8-bit grayscale (Image → Type → 8-bit). Next, the image was converted to an “invert” image to ensure high pixel values correspond to the dense (protein-rich) bands while low values for the background (Edit → Invert). Lanes were defined using the gel analysis function (Analyze → Gels), in which a rectangular region of interest of fixed dimensions was drawn around the band of the target protein in the first lane along with the entire lane respectively and then applied uniformly to all remaining lanes to ensure identical measurement areas across samples. The intensity corresponding to the region of interest was estimated using the “Measure” drop-down option under the “Analyze” tool to obtain the integrated band density along with the mean gray value. The resultant values were plotted in the form of bar graphs using origin pro.

### Hydrogen Deuterium Exchange Mass Spectrometry

Deuterium labeling experiments were performed for different biological states namely-purified protein, cellular lysate, and intact cells. All the labelling reactions were performed under identical experimental conditions to allow for direct comparisons between the different states. To monitor the binding effects of BI2852 on the KRAS in a cellular lysate, cell lysate containing overexpressed KRAS protein or purified KRAS was incubated with BI2852 at 1:2 molar ratio and was incubated at room temperature (23 °C) for 30 minutes prior to deuterium labeling. HDX of the cell lysate, cell suspension and protein alone was initiated with a 9-fold dilution into D_2_O labelling buffer (20 mM HEPES, 150mM NaCl 1 mM TCEP, pH 8.0 in 99.9% D_2_O). After each labelling time (10 seconds, 1 minute, 10-minutes, 30 minutes, and 1 hour) at 23 °C, the labelling reaction was quenched with ice-cold buffer containing 5M Guanidium HCl, 0.8% formic acid, 2.5% DDM, 0.8M TCEP, 2mg Aspergillus protease, pH 2.3 and was snap frozen in liquid nitrogen followed by LC-MS analysis. Control samples were prepared using the same method as above except using a buffer containing 20 mM HEPES, 150mM NaCl 1 mM TCEP, pH 8.0 in H_2_O. All deuterium-labeling experiments were performed in duplicate (for the lysate and purified protein) and triplicate (for intact cells) respectively. The quenched samples were digested online (15° C) using an in-house pepsin column. The resultant peptides were desalted for 3 minutes at a flow rate of 100 μl/min using a VanGuard Pre-Column trap (2.1 mm × 5 mm, ACQUITY UPLC BEH C18, 1.7 μm). Elution of peptides from the trap was achieved using a 5-35% acetonitrile gradient with 0.1% formic acid over 20 minutes at a flow rate of 100 μl/min and peptides were separated using an ACQUITY UPLC HSS T3 column (1.8 μm, 1.0 mm × 50 mm). The cooling chamber of the HDX system was maintained at 0.0 ± 0.1°C throughout the measurements.

#### Mass spectrometry analysis

Mass spectra were acquired using a Waters SELECT SERIES Cyclic IM QTOF mass spectrometer. Spectra were acquired between 50 and 2000 *m*/*z* with capillary voltage 3.0 kV, sample cone 40 V, and transfer CE ramping from 20 to 40 V. Single-pass cIM was performed with a 10 ms injection, 3 ms separation, and 34 ms ejection/acquire sequence with a TW static height of 23 V and an analog-to-digital converter (ADC) start delay of 13 ms and using one pushes per bin. The error in determining the average deuterium incorporation for each peptide was at or below ±0.25 Da, based on deuterated peptide standards.

Peptide identification was performed in Protein Lynx Global Server (PLGS) v3.0.1 (Waters Corporation) using replicates of undeuterated control samples. Peptide masses were identified from searches using non-specific cleavage of custom database containing the sequences of both His_6_-tagged and untagged KRAS (Uniprot ID: P01116), SOSCAT (Uniprot ID: Q07889), GSK3β (Uniprot ID: P49841) and GK (Uniprot ID: P35557) along with 130 bacterial proteins that were identified from a tryptic digest of lysate derived from *Ecoli* BL21 DE3 cells in a separate experiment. The raw MS data were imported into DynamX 3.1.1 (Waters corporation) and filtered implementing a minimum product per amino acid cut-off of 0.1, at least 1 consecutive product ions. Those peptides meeting initial filtering criteria were further processed automatically by DynamX followed by manual inspection of all spectra. The relative amount of deuterium in each peptide was determined by subtracting the centroid mass of the undeuterated form of each peptide from the deuterated form, at each timepoint for each condition. All uptake and difference plots, found in the Figures and Supplementary data were generated using these deuterium incorporation values, and the complete dataset is provided in Supplementary Data excel file. Deuterium levels were not corrected for back exchange and thus reported as relative (9). All HDX-MS data have been deposited to the ProteomeX-change Consortium via the PRIDE (51) partner repository with the dataset identifiers PXD087555 and 10.6019/PXD087555 as well as PXD087155 and 10.6019/PXD087155.

## Supporting information

Supplemental Datafile

